# CRISPR-TAPE: protein-centric CRISPR guide design for targeted proteome engineering

**DOI:** 10.1101/2020.02.26.965889

**Authors:** DP. Anderson, HJ. Benns, EW. Tate, MA Child

**Author notes:** These authors contributed equally: Daniel P Anderson and Henry J Benns.

## Abstract

Rational molecular engineering of proteins with CRISPR-based approaches is challenged by the gene-centric nature of gRNA design tools. To address this, we have developed CRISPR-TAPE, a protein-centric gRNA design algorithm that allows users to target specific residues, or amino acid types within proteins. gRNA outputs can be customized to support maximal efficacy of homology-directed repair for engineering purposes, removing time consuming post-hoc curation, simplifying gRNA outputs, and reducing CPU times.

## Main Text

Functional genomics has been revolutionised by the discovery of the CRISPR prokaryotic defence system^1,2^ and its conversion into an effective and efficient mechanism for genome engineering^3-5^. An abundance of bioinformatic tools are available for the identification and selection of unique CRISPR guide RNA (gRNA) sequences required for nuclease targeting of individual genes^6,7^, but search algorithms have remained gene-centric (Fig. S1). This has limited the wider exploitation of these technologies by protein- and proteome-engineers, and researchers seeking to modify specific amino acids or protein sequences such as catalytic residues within enzyme active sites.

At present, gRNAs targeting protein-coding regions within a genomic locus are anonymous within total gRNA outputs from existing design tools that non-specifically target the entire input region of DNA (Fig. S2). These gRNA lists subsequently require extensive time-consuming manual curation to identify those targeting specific protein regions of interest. The principle challenge encountered by existing design algorithms is the non-linear correlation of genomic sequence (intronic and exonic) with protein coding sequence (spliced exonic). This often requires users to manually distinguish between intron-exon sequences and pair target amino acid/s with proximal gRNAs. The gene-centric focus of existing algorithms has inadvertently led to an absence of protein design considerations. These include distance of the nuclease cut-site from a protein feature of interest for efficient homology-directed repair (HDR)^8^, and an inability to query gRNAs targeting specific amino acids or positions of interest such as sites of post-translational modification (PTM).

To address this, we have developed CRISPR-TAPE, a protein-centric CRISPR gRNA design algorithm for Targeted Proteome Engineering (Fig. 1a, S3 and online methods). CRISPR-TAPE is run as a freely available stand-alone python script, and custom executable application (www.laboratorychild.com/crispr-tape) (Fig. 1b and supplemental note). The CRISPR-TAPE algorithm is based upon a multidimensional matrix mapping strategy, where the coding sequence is split into codons and these translated to a protein sequence. The corresponding codon positions relative to the genomic sequence are stored for each residue. This numerical indexing enables the algorithm to account for complex intron-exon gene structures and supports protein-focused gRNA searches. Users query amino acid types (e.g. cysteines) or individual amino acids at specific positions of interest within their input protein sequence (e.g. cysteine 106) facilitating direct interrogation of protein domains, catalytic motifs, and PTMs. The reduced complexity of the search space greatly reduces computational burden and central processing unit (CPU) times for gRNA identification; applied to the ∼80 Mb genome of the model eukaryotic pathogen *Toxoplasma gondii*, for a single specified amino acid within an input protein coding sequence, guides directing Cas9 to within 30 nucleotides of the residue for optimal HDR are typically output in less than five seconds (Fig. S4a). Although comparisons with existing tools are inherently unfair (due to fundamental differences in source codes), identical searches using existing gene-centric tools require a processing time in the order of minutes. For a better comparison, we benchmarked CRISPR-TAPE performance against our source code; we deployed our code for non-directed guide identification within genomic loci (i.e. traditional total guide output for genomic loci). Compared with our own source code deployed for total guide identification, CPU times required by CRISPR-TAPE for searches are greatly reduced. gRNAs for position-specific queries are processed at an average rate of ∼330 nucleotides per second (Fig. S4a). For type-specific queries, using leucine as a linearly distributed amino acid in our test gene set (Fig. S4b), gRNAs are processed at an average rate of ∼27 nucleotides per second (Fig. S4a). These rates are approximately 40- and 3.2-fold faster than basic gRNA identification for the position-specific and type-specific search modes respectively.

**Figure 1:**
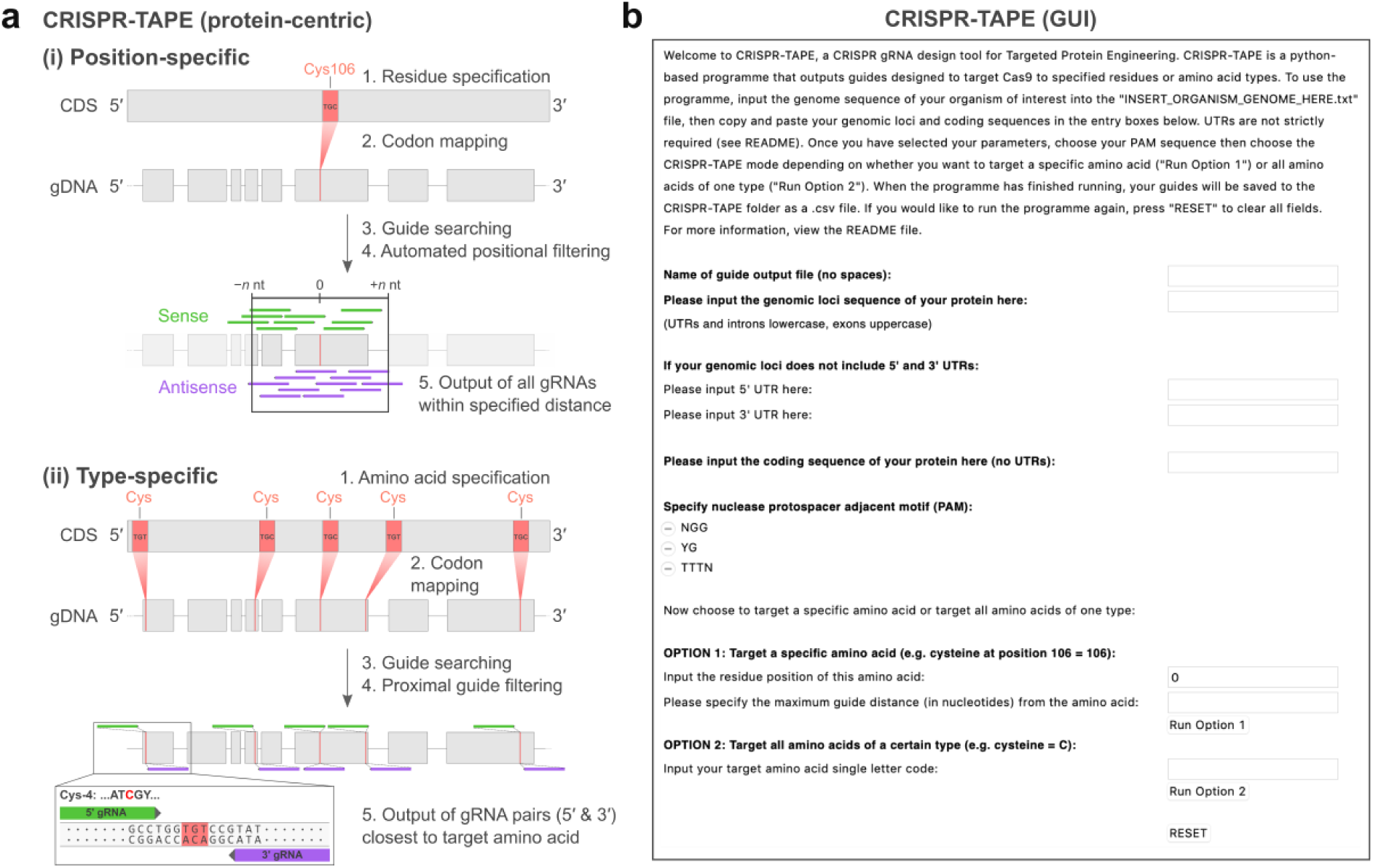
Automated protein-centric identification of CRISPR gRNAs using CRISPR-TAPE. (**a**) Workflow schematics of position-specific (i) and type-specific (ii) functions of CRISPR-TAPE. Following selection of a specific residue position (i) or amino acid type (ii; 1), the corresponding codon(s) from the target protein’s coding sequence (CDS) are mapped onto its genomic DNA (gDNA) locus (2). A selected PAM sequence is then queried against the gDNA to identify guides (3) and filtered according to a user-defined cut site distance from the specified residue position (i) or by the two most proximal gRNAs that target up- and downstream of each protein sequence representation of an amino acid type (ii; 4). (**b**) Graphic user interface (GUI) of CRISPR-TAPE.

CRISPR-TAPE reduces output gRNA complexity as guide sequences are automatically curated. gRNA outputs are provided in relation to the specified amino acid or amino acid type within the genomic locus and distributed according to distance of the nuclease cut site from the specified amino acid/s to support efficient HDR strategies (Fig. 2). gRNAs therefore require no further manual curation to correlate genome targeting with the protein coding sequence (unlike traditional gene-centric gRNA outputs, Fig. S2). While it is challenging to quantify time-saving performance improvements when comparing automated to manual curation (as it involves highly variable parameters such as user experience and background knowledge), the benefit of automated curation of gRNAs when using CRISPR-TAPE is expected to be substantial. Distinct from gene-centric search tools, CRISPR-TAPE prioritizes proximity of the nuclease cut-site to a specified amino acid type/position over all other guide selection and scoring criteria.

**Figure 2:**
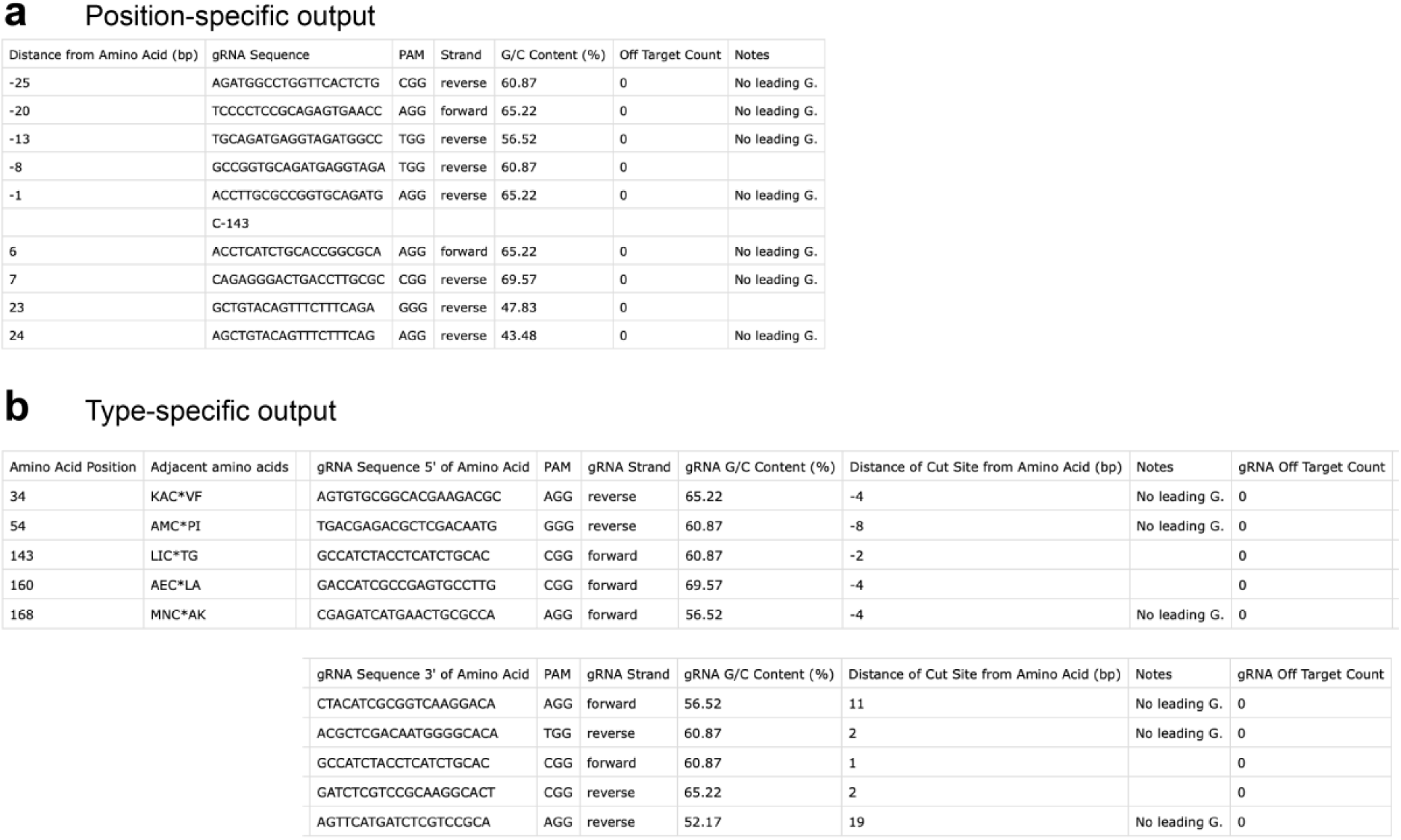
Output files generated by CRISPR-TAPE using the latest *Toxoplasma gondii* GT1 genome release (ToxoDB-46), and querying *Toxoplasma* gene ID TGGT1_242330 with (**a**) the “Position-specific” function, identifying gRNAs within 30 nucleotides of a target cysteine at position 143, and (**b**) the “Type-specific” function, with gRNA pairs provided for all cysteines present in the CDS.

CRISPR-TAPE performance was then assessed for a potential genome-scale protein engineering application; we identified a well-characterised dataset for the malaria parasite *Plasmodium falciparum*, where a known PTM (*S*-palmitoylation) has been globally profiled but site-specific information regarding the modified amino acid (cysteine) is absent^11^. Such a dataset is suitable for the CRISPR-TAPE “type-specific” function, where pairs of gRNAs proximal to a user defined amino acid type are output simultaneously (Fig. 2b). We focused on 55 putatively palmitoylated proteins the authors of this work identified as being representative of common and uncommon palmitoyl protein classes^11^. We first used a traditional gene-centric search algorithm^10^ to identify all gRNAs present within the input gene sequences. CPU time required for the initial gRNA search was on average 25 seconds per gene. In our experience, manual curation time required to identify pairs of gRNAs targeting individual amino acids takes a minimum of five minutes/residue. Considering that the average number of cysteines/gene in this dataset is 11 and including the average CPU time, this can be extrapolated to ∼55 minutes per gene. We then applied CRISPR-TAPE to the same dataset, using the “type-specific” option to identify pairs of gRNAs in proximal positions to all potential sites of *S*-palmitoylation (i.e. all cysteine residues). The average processing time for each protein was six seconds, with no downstream curation of gRNA outputs required. Applied in this way and compared with a traditional gene-centric gRNA design tool, CRISPR-TAPE was at least 300 times faster per cysteine being targeted. These data are summarized in Table S1.

The profiling of amino acid reactivity within proteins is a rapidly expanding field, with this chemical proteomic strategy having been successfully applied to profile the reactivity of serine^12^, cysteine^13^, lysine^14^, histidine^15^, tyrosine^16^, and methionine^17^ residues. A method to efficiently prioritize these reactive sites according to their contribution to protein function would optimize pipelines for target-based screening platforms. Such a method could be supported by CRISPR-based site-directed mutagenesis via homology-directed repair. To test the performance of our algorithm applied to specific sites of interest, we selected a subset of 12 ligandable sites on 10 proteins from a recent reactive lysine profiling dataset within the human cancer cell proteomes of MDA-MB-231, Ramos and Jurkat cells^14^. We focused on kinases as targets of proven therapeutic value, and sought to identify a panel of gRNAs that would direct Cas9 to within 30 nucleotides of a target lysine (to support efficient HDR for mutational interrogation of its function). CPU times for CRISPR-TAPE were on average double that of the online gene-centric tool CHOPCHOP^18^. This likely reflects the power of the consortia-funded dataservers that support these community tools, and makes direct comparison of processing speeds for different algorithms inherently unequal. On average, CRISPR-TAPE identified 10 gRNAs/reactive lysine within the defined 30 nt window upstream and downstream of the residue codon, with these gRNA panels auto-curated and immediately ready for downstream applications. A conservative estimation for the manual curation of tightly targeted gRNA panels from dense gRNA output arrays provided by gene-centric tools would be a minimum of five minutes/gRNA. For the identical panel of 10 gRNAs/reactive lysine, this can be extrapolated to an expected total manual curation time of 50 minutes/lysine. Accounting for differences in raw processing speed this makes CRISPR-TAPE at least 10 times faster than gene-centric tools used in this way, and these data are summarized in Table S2.

CRISPR-TAPE source code is freely available (github.com/LaboratoryChild/CRISPR-TAPE), organism-adaptable and includes standard guide-design features such as off-target scoring and identification of guide sequences predicted to be ineffective^3,7,9^. The code can also be expanded to include amino acid motif-based proteome engineering strategies and batch processing. We anticipate that CRISPR-TAPE will support existing gene-centric tools and empower the proteome engineering community. It will accelerate the application of CRISPR-based methods for targeted protein modification, *in vivo* protein evolution, and amino acid prioritization in drug discovery.

## Acknowledgements

This work was supported by grants BB/M011178/1 from the BBSRC (to H.J.B., E.W.T., and M.A.C.) and 202553/Z/16/Z from the Wellcome Trust & Royal Society (to M.A.C.). We would also like to thank Professor Mike Sternberg, and Dr Ellen McDonagh for critical reading of the manuscript, and beta-testers in the Child and Tate laboratories.

## Online Computational Methods (see also Fig. S3)

### User Input

CRISPR-TAPE requires several user inputs to function correctly. Once CRISPR-TAPE has loaded, user inputs are added via entry boxes in the graphical user interface (GUI) window. Both running modes (custom application or raw Python script) require the organism genome to be accessible for successful gRNA off-target site identification. This is done by downloading the genome of the organism of interest, placing the genome file within the specified location (see below), and then renaming it to “INSERT_ORGANISM_GENOME_HERE.txt” (and replacing the existing file with that name). Genome files can be downloaded in FASTA format (.fa) and directly renamed. The genome .txt file must be located within the same directory as the CRISPR-TAPE executable application (or Python script if using command line) for the programme to recognise and import the genome of interest when running.

Within the GUI window, the user begins by specifying a filename for the guide RNA (gRNA) output table. Users then specify input the genomic loci sequence of the protein of interest (5’ to 3’ orientation), with introns and UTRs in lowercase and exons in uppercase. If UTRs are not included, users can also include 100 bases (or longer) upstream and downstream of the genomic loci to enable identification of gRNAs targeting amino acids in close proximity to the protein N- and C-termini, respectively. While this is not strictly required and these input boxes can be left empty, exclusion of flanking sequences may limit the number of gRNAs identified at these sites. CRISPR-TAPE currently supports ‘NGG’, ‘YG’ and ‘TTTN’ protospacer adjacent motifs (PAMs) and these can be selected through a checkbox. The open source availability and modularity of the CRISPR-TAPE base code allows for easy incorporation of additional PAM sequences (see the download-associated README file and Supplemental Note 2 for detailed information). Users then choose to target an amino acid at a specific position by inputting the position of the residue within the protein sequence (OPTION 1), or target all amino acids of a certain type by inputting the amino acid single letter code (OPTION 2). Within OPTION 1 the user may also specify the maximum base distance of guides from the residue of interest to limit the range of outputted gRNAs. If no distance is specified, all guides within the input locus are identified.

### Pre-processing

The pre-processing stage is required to import and store the genome of the organism of interest from the “INSERT_ORGANISM_GENOME_HERE.txt” and perform some basic manipulations of the user inputted sequences. The inputted genomic loci and stored genome are reverse complemented for reverse strand guide identification and off-target counting, respectively. The uppercase bases within the genomic loci are recognized and converted into a coding sequence (CDS). This concatenated exon sequence is then aligned to the user inputted CDS. If these are not identical, gRNAs will not be generated. This launches an error pop-up informing the user that that the intron-exon structure of the gene in relation to the coding sequence do not match and should be checked. The positions of exonic bases relative to the genomic loci are then stored in a 1D array for matrix generation in the next stage. The inputted CDS is translated into an amino acid sequence and the matrix position of each residue is stored.

## CRISPR-TAPE

### OPTION 1 and 2 shared processes

A multidimensional matrix is generated to compile information for each amino acid within the input CDS. This matrix consists of the position of each residue in the protein sequence, the specific codon that codes for that amino acid, and the position of each of the corresponding codon bases within the inputted genomic loci. The programme searches for the positions of the user-specified PAM on both the forward and reverse strands of the genomic locus and outputs the position of the base immediately 5’ of the nuclease cut site. This position is later used to determine an initial crude distance value between the cut site and the base 5’ of the codon. This positional information is used to generate forward and reverse strand lists of all potential gRNAs within the inputted genomic loci, arranged by position. gRNA sequences and associated nuclease cut site positions are converted to a dataframe and G/C percentages of gRNA sequences calculated.

The matrix position of the 5’ or 3’ base in the codon triplet is used to determine which base 5’ of the gRNA(s) cut site position is in closest proximity to the user specified residue(s) within the genomic loci, and this used to identify the corresponding gRNA sequences 5’ and 3’ to the target. When gRNAs are initially identified, their position is indexed as the base 5’ of the nuclease cut site regardless which strand they are on in relation to the inputted genomic locus. A subsequent function then corrects this distance accounting for the strand; The distance between the 5’ or 3’ base of the nuclease cut site and 5’ or 3’ base of the codon is calculated and appended to the gRNA dataframe(s). “Leading G” and “poly-T” information of gRNA sequences is also determined and appended to the dataframe. Off-targets are searched and counted for each gRNA sequence in the inputted organism genome and its reverse complement. The tool does not provide a score for gRNA outputs.

### OPTION 1 specific

Guides located beyond the user-specified distance are removed from the dataframe. The gRNA dataframe is then split (based on positional information) into gRNAs 5’ and 3’ of the residue. The dataframe of 5’ gRNAs is arranged by decreasing distance between cut site and amino acid, and the dataframe of 3’ gRNAs arranged by increasing distance.

### OPTION 2 specific

gRNAs with > 75% G/C content and associated cut site positions are removed from the dataframe.

### Output

The list of the gRNAs generated by CRISPR-TAPE are outputted within the CRISPR-TAPE home directory to a “.csv” file specified by the user.

**The output of OPTION 1 consists of:**

1. **gRNA Sequence**: The gRNA sequence identified by the programme.
2. **PAM**: The specific protospacer adjacent motif immediately adjacent to the gRNA
3. **Strand**: The orientation of the DNA strand the gRNA targets relative to the sense of the inputted genomic loci.
4. **G/C Content**: The percentage of the gRNA sequence consisting of “G” and “C” bases.
5. **Distance from aa (bp)**:
  - If positive and “Strand” = forward: gRNA is upstream of the amino acid, the distance is measured from the base on the right-hand side of the nuclease cut site to the base on the left hand-side of the codon (5’to 3’).
  - If negative and “Strand” = forward: gRNA is downstream of the amino acid, the distance is measured from the base on the left-hand side of the nuclease cut site to the base on the right hand-side of the codon (5’to 3’).
  - If positive and “Strand” = reverse: gRNA is upstream of the amino acid, the distance is measured from the base on the left-hand side of the nuclease cut site to the base on the left hand-side of the codon (5’to 3’)
  - If negative and “Strand” = reverse strand: gRNA is downstream of the amino acid, the distance is measured from the base on the right-hand side of the nuclease cut site to the base on the right hand-side of the codon (5’to 3’).

1. **Notes:** Does the gRNA contain a poly-T sequence indicated by a tandem of 4 or more Ts? Does the gRNA have a leading G at position 1 in the gRNA? Is the G/C content over 75%?
2. **Off-target Count**: The number of off-target sites the gRNA may target.

**The output of OPTION 2 consists of**

1. **Amino Acid Position**: The position of the amino acid within the amino acid sequence.
2. **Adjacent amino acids**: The 4 amino acids immediately surrounding the residue being targeted. The target residue is indicated by “*”
3. **5’ gRNA Sequence**: The sequence of the gRNA closest in proximity upstream of the amino acid.
4. **3’ gRNA Sequence**: The sequence of the gRNA closest in proximity downstream of the amino acid.
5. **PAM**: The specific protospacer adjacent motif immediately adjacent to the 5’ or 3’ gRNA.
6. **Strand**: The orientation of the DNA strand the 5’ or 3’ gRNA targets relative to the sense of the inputted genomic loci.
7. **G/C Content**: The percentage of the 5’ or 3’ gRNA sequence consisting of “G” and “C” bases.
8. **Distance of cut site from Amino Acid (bp)**:
  - If “Strand” = forward and the gRNA is 5’ of the residue: The distance is measured from the base on the right-hand side of the nuclease cut site to the base on the left-hand side of the codon (5’to 3’).
  - If “Strand” = forward and the gRNA is 3’ of the residue: The distance is measured from the base on the left-hand side of the nuclease cut site to the base on the right-hand side of the codon (5’to 3’).
  - If “Strand” = reverse and the gRNA is 5’ of the residue: The distance is measured from the base on the left-hand side of the nuclease cut site to the base on the left-hand side of the codon (5’to 3’).
  - If “Strand” = reverse and the gRNA is 3’ of the residue: The distance is measured from the base on the right-hand side of the nuclease cut site to the base on the right-hand side of the codon (5’to 3’).
9. **Notes**: Does the 5’ or 3’ gRNA contain a poly-T sequence indicated by a tandem of 4 or more Ts? Does the gRNA have a leading G at position 1 in the gRNA? Is the G/C content over 75%?
10. **Off-target Count**: The number of off-target sites the 5’ and 3’ gRNA may target.

## Supplemental Material

**Figure S1:**
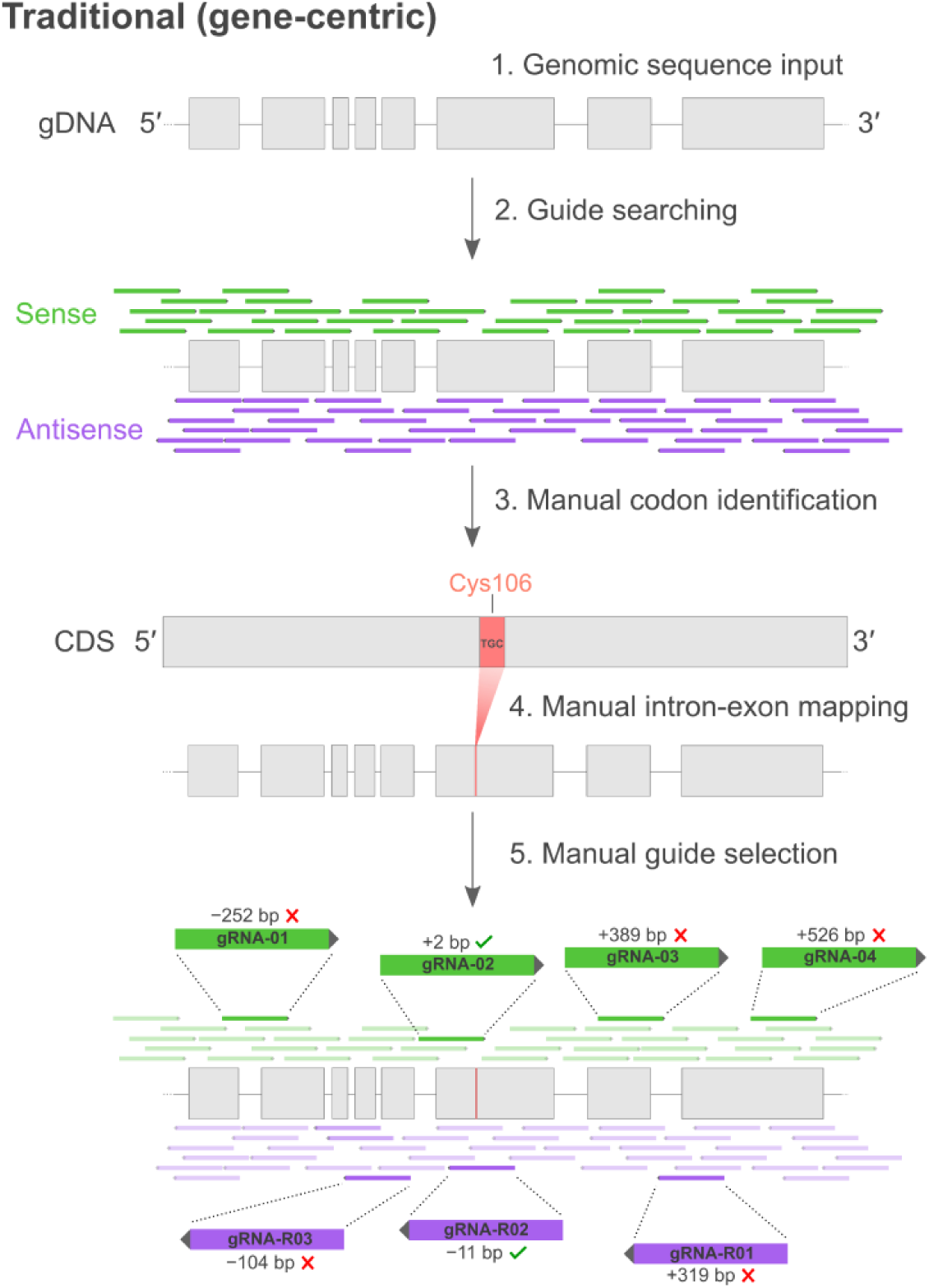
Traditional gene-centric approach for amino acid-targeted CRISPR gRNA design. Users input the genomic DNA (gDNA) locus of their target protein into an existing gene-centric gRNA design tool (1). Gene-centric algorithms typically query chosen PAM sequences against the entire genomic locus without subsequent positional filtering, resulting in a complex output of large numbers of gRNAs (2, and Fig. S2). The codon positions of the target amino acid(s) are then manually identified in the protein coding sequence (CDS; 3) and mapped onto its associated gDNA sequence, for users to be able to account for complex intron-exon gene structures (4). From the extensive list of gRNAs generated in 2, users must initiate time-consuming manual curation to identify gRNAs with a nuclease cut site in close proximity to residues on both sense and antisense gDNA strands (5).

**Figure S2:**
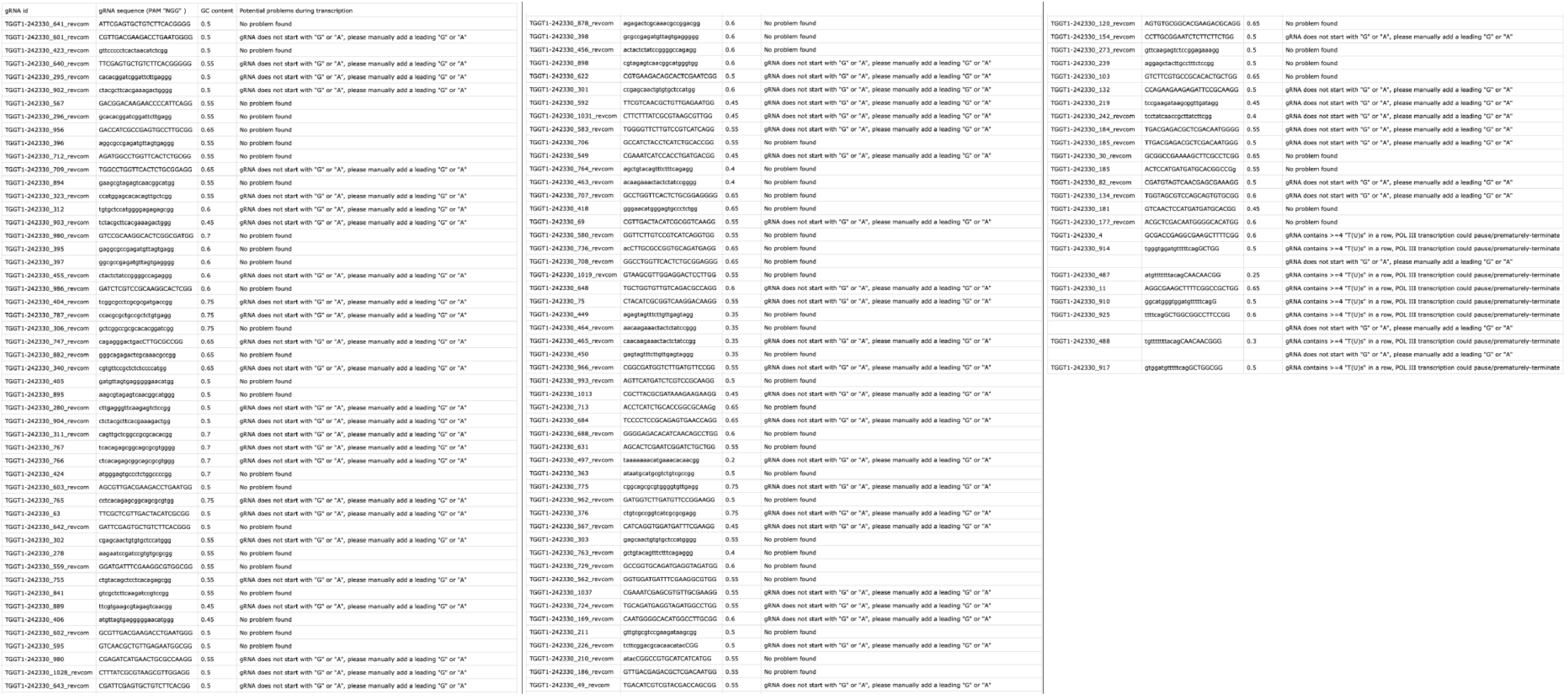
Example gRNA output generated using the Eukaryotic Pathogen CRISPR guide RNA/DNA Design Tool_10_ using a server-hosted *Toxoplasma gondii* GT1 genome release (ToxoDB-32), and querying *Toxoplasma* gene ID TGGT1_242330. Data presented is a simplified list of full dataset generated.

**Figure S3:**
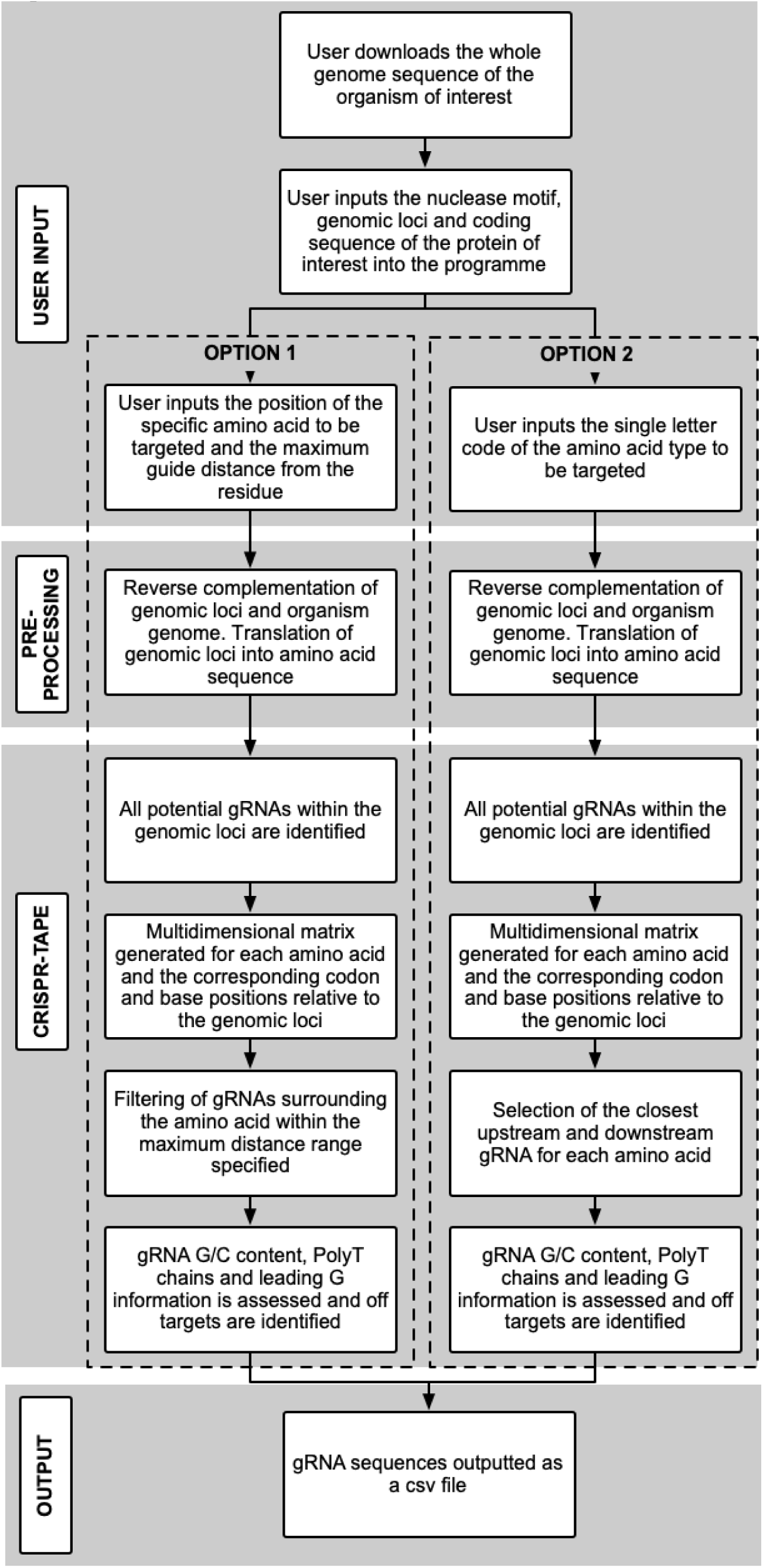
Flowchart schematic of CRISPR-TAPE algorithm.

**Figure S4:**
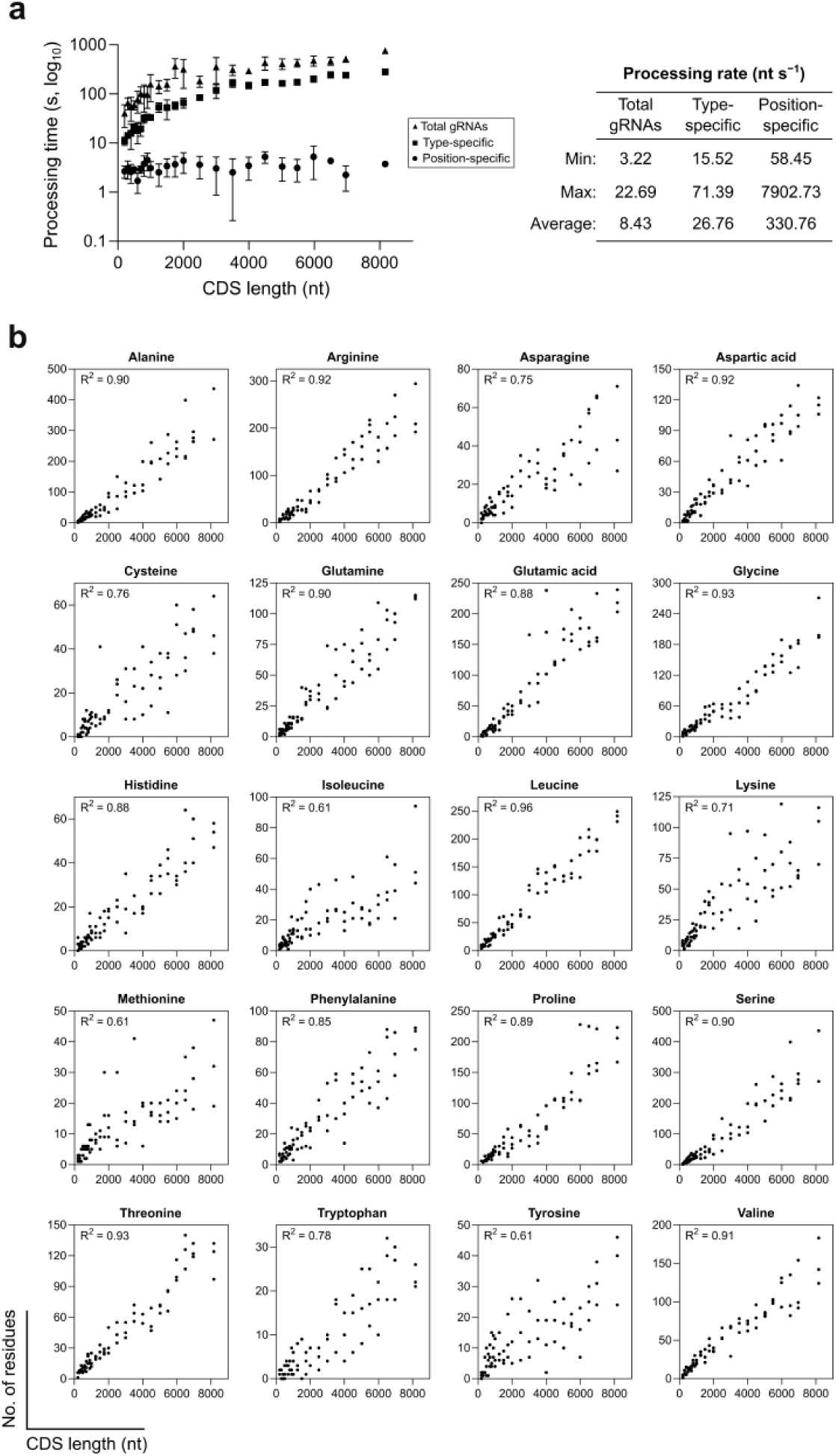
CRISPR-TAPE performance comparisons. (**a**) *Left:* comparison of processing speeds of CRISPR-TAPE versus total gRNA output for genes from the latest *Toxoplasma gondii* GT1 genome release (ToxoDB-46). Mean (±SD) gRNA output processing times are displayed for three genes of various CDS lengths (n=3) with ‘*NGG*’ specified as the target protospacer adjacent motif (PAM). For the position-specific function, gRNAs were queried within a distance of 30 nucleotides (nt) from methionine at position 1. For the type-specific function, leucine was specified as the target amino acid. *Right:* minimum, maximum and mean processing rates for each function across the same dataset. (**b**) Linear regression analyses of amino acid frequency and CDS length across the gene panel analysed in A. Coefficients of determination (R_2_) are indicated for each amino acid type.

**Table S1:**

| **Palmitoylated cysteines in *P. falciparum*** |  |  |  |
| --- | --- | --- | --- |
|  | EuPaGDT | CRISPR-TAPE<br>(type-specific) | Relative performance<br>of CRISPR-TAPE |
| No. of genes: | 55 | 55 | N/A |
| No. of cysteines: | 584 | 584 | N/A |
| Total CPU time (s)*: | 1,362 | 348 | ~4× faster |
| Total gRNA curation time (s): | 175,200 | 0 | > 300× faster/cysteine |
| **Total time (h):** | **49.1** | **0.1** | **~500× faster** |

\*gRNAs processed in batch (EuPaGDT) or individually (CRISPR-TAPE)

**Table S2:**

| **Ligandable lysines in *H. sapiens*** |  |  |  |
| --- | --- | --- | --- |
|  | CHOPCHOP | CRISPR-TAPE<br>(position-specific) | Relative performance<br>of CRISPR-TAPE |
| No. of genes: | 10 | 10 | N/A |
| No. of ligandable lysines: | 12 | 12 | N/A |
| Total CPU time (s)*: | 2,002 | 3,980 | ~2× slower |
| Total gRNA curation time (s)**: | 36,000 | 0 | > 3,000× faster/lysine |
| **Total time (h):** | **10.6** | **1.1** | **~10× faster** |

\*gRNAs processed individually \*\*Curation time of 300 s/gRNA for an average of 10 gRNAs identified/lysine

**Supplemental Note: Use of CRISPR-TAPE (application-associated README document)**

### Overview

CRISPR-TAPE is a python-based CRISPR design tool for TArgeted Protein Engineering, available as a standalone application for Windows and macOS operating systems. The CRISPR-TAPE source code is open source and their modularity allows for easy incorporation of additional functions (github.com/LaboratoryChild/CRISPR-TAPE). Users may achieve enhanced CPU times using the raw python scripts and this is recommended for computationally intensive guide RNA (gRNA) generation (e.g. for genomes larger than 1 giga-base). The standard CRISPR-TAPE application incorporates a custom graphic user interface (GUI) and is recommended for most users working with genomes less than one giga-base in size.

### Motivation

Existing CRIPSR gRNA design tools non-specifically target the entire input region of DNA. Current gene-centric design algorithms fail to consider protein-focused users. CRISPR-TAPE has been developed to reduce the substantial time burden associated with manual curation of gRNA libraries, and make CRISPR-based protein modification strategies more accessible to the protein engineering community.

### Current version

CRISPR-TAPE version 1.0.0

### System requirements

Systems running CRISPR-TAPE must meet the following requirements:

- macOS version 10.9.5 or later, 4 GB+ RAM
- Windows 7 or later, 4 GB+ RAM

For target organisms with genomes greater than one giga-base in size, it is recommended systems have at least 32 GB RAM to prevent application crashes, with <u>64 GB+ RAM recommended for genomes greater than three giga-bases in size (e.g.</u> *<u>Homo sapiens</u>*).

### Installation

The following provides and overview for the installation process. A detailed walk-through with example sequences is provided in the “**Walk-through and test example sequences**” section below.

- CRISPR-TAPE for Windows and macOS is available from via http://www.laboratorychild.com/crispr-tape
  ⇒For Windows-based systems, download and unzip the CRISPR-TAPE.exe file and place it in a directory of your choice.
  ⇒On macOS systems, the CRISPR-TAPE.dmg should be downloaded, mounted, and the CRISPR-TAPE application file dragged and dropped into the Application alias folder when prompted.
- For successful gRNA identification, the genome of the organism of interest must be located within a single .txt file titled “INSERT_ORGANISM_GENOME_HERE.txt”. Genome files are typically available for download in FASTA format, and can be directly renamed “INSERT_ORGANISM_GENOME_HERE.txt”. This renamed file can then be moved to the appropriate file directory (detailed above), replacing the exiting file in that location.
  ⇒On Windows, this file must be located within the same directory as the CRISPR-TAPE.exe file.
  ⇒On macOS the file is located within the CRISPR-TAPE.app package via the following path: “./CRISPR-TAPE.app/Contents/Resources”, accessible by right clicking the application and pressing “Show Package Contents” => subdirectory “Contents” => subdirectory “Resources”.

These respective Windows and macOS directories are also where gRNA .csv files are outputted to once the programme has run successfully.

### Input

Within the CRISPR-TAPE GUI, users input the filename of the outputted .csv file and the genomic loci and coding sequences of the protein of interest. Users then specify the protospacer adjacent motif (PAM) sequence via a checkbox. To target a specific residue, users input the numerical position of the residue within the protein sequence and the maximum distance for gRNA identification if desired, then choose “Run Option 1”. To target all amino acids of one type, input the single letter code of the residue of interest and choose “Run Option 2”.

### Output

The list of the gRNAs generated by CRISPR-TAPE are outputted to a “.csv” file specified by the user within the CRISPR-TAPE home directory on Windows and within “./CRISPR-TAPE.app/Contents/Resources” on macOS.

The dataframe outputted by Specific_function consists of:

1. gRNA Sequence: The gRNA sequence identified by the programme.
2. PAM: The specific protospacer adjacent motif immediately adjacent to the gRNA
3. Strand: The orientation of the DNA strand the gRNA targets relative to the sense of the inputted genomic loci.
4. G/C Content: The percentage of the gRNA sequence consisting of “G” and “C” bases.
5. Distance from aa (bp):
  - If positive and “Strand” = forward: gRNA is upstream of the amino acid, the distance is measured from the base on the right-hand side of the nuclease cut site to the base on the left-hand side of the codon (5’to 3’).
  - If negative and “Strand” = forward: gRNA is downstream of the amino acid, the distance is measured from the base on the left-hand side of the nuclease cut site to the base on the right-hand side of the codon (5’to 3’).
  - If positive and “Strand” = reverse: gRNA is upstream of the amino acid, the distance is measured from the base on the left-hand side of the nuclease cut site to the base on the left-hand side of the codon (5’to 3’)
  - If negative and “Strand” = reverse strand: gRNA is downstream of the amino acid, the distance is measured from the base on the right-hand side of the nuclease cut site to the base on the right-hand side of the codon (5’to 3’).
6. Notes: Does the gRNA contain a poly-T sequence indicated by a tandem of 4 or more Ts? Does the gRNA have a leading G at position 1 in the gRNA? Is the G/C content over 75%?
7. Off-target Count: The number of off-target sites the gRNA may target.

The dataframe outputted by General_function consists of:

1. Amino Acid Position: The position of the amino acid within the amino acid sequence.
2. Adjacent amino acids: The 4 amino acids immediately surrounding the residue being targeted. The target residue is indicated by “*”
3. 5’ gRNA Sequence: The sequence of the gRNA closest in proximity upstream of the amino acid.
4. 3’ gRNA Sequence: The sequence of the gRNA closest in proximity downstream of the amino acid.
5. PAM: The specific protospacer adjacent motif immediately adjacent to the 5’ or 3’ gRNA.
6. Strand: The orientation of the DNA strand the 5’ or 3’ gRNA targets relative to the sense of the inputted genomic loci.
7. G/C Content: The percentage of the 5’ or 3’ gRNA sequence consisting of “G” and “C” bases.
8. Distance of cut site from Amino Acid (bp):
  - If “Strand” = forward and the gRNA is 5’ of the residue: The distance is measured from the base on the right-hand side of the nuclease cut site to the base on the left-hand side of the codon (5’to 3’).
  - If “Strand” = forward and the gRNA is 3’ of the residue: The distance is measured from the base on the left-hand side of the nuclease cut site to the base on the right-hand side of the codon (5’to 3’).
  - If “Strand” = reverse and the gRNA is 5’ of the residue: The distance is measured from the base on the left-hand side of the nuclease cut site to the base on the left-hand side of the codon (5’to 3’).
  - If “Strand” = reverse and the gRNA is 3’ of the residue: The distance is measured from the base on the right-hand side of the nuclease cut site to the base on the right-hand side of the codon (5’to 3’).
9. Notes: Does the 5’ or 3’ gRNA contain a poly-T sequence indicated by a tandem of 4 or more Ts? Does the gRNA have a leading G at position 1 in the gRNA? Is the G/C content over 75%?
10. Off-target Count: The number of off-target sites the 5’ and 3’ gRNA may target.

### Walk-through and test example sequences

Here, we provide a basic walk-though for new users, including test input sequences for the *Toxoplasma gondii* gene TGGT1_242330. The outputs from user tests can be directly compared against those in Figure 2 in the associated manuscript (see **Citation** below).

### Step

1. CRISPR-TAPE comes preloaded with the *Toxoplasma gondii* GT1 genome (release 46). The genome file was downloaded in FASTA format and directly renamed “INSERT_ORGANISM_GENOME_HERE.txt”. This file was then moved to the appropriate file directory (detailed in **Installation** above), replacing the existing file in that location.
2. Following installation launch the CRISPR-TAPE application.
3. Choose a file name for the gRNA outputs, and type in the input box: “**Name of guide output file (no spaces)**”. For this example, name the file “**test**”.
4. Copy-paste the following sequence into the “**Please input the genomic loci sequence of your protein here**” box. Note that the gene intron-exon structure is distinguished though the use of lower- and uppercase, respectively. The gene sequence used for this test example does not include UTRs: ATGGCGACCGAGGCGAAGCTTTTCGGCCGCTGGTCGTACGACGATGTCAACGTCAGCGACCTTTCGCTCGTTGACTACATCGCGGTCAAGGACAAGGCCTGCGTCTTCGTGCCGCACACTGCTGGACGCTACCAGAAGAAGAGATTCCGCAAGGCCATGTGCCCCATTGTCGAGCGTCTCGTCAACTCCATGATGATGCACGGCCGgtatgttgtgcgtccgaagataagcggttgataggagctacttgcctttctccggagactcttgaaccctcaagaatccgatccgtgtgcgcggccgagcaactgtgtgctccatggggagagagcggaacacgcgtattcatgtatctgtgtattataatgcatgcgtctgtcgccggtcatcgcgcgaggcgccgagatgttagtgagggggaacatgggagtgccctctggccccggatagagtagtttcttgttgagtagggatccgttgtgtttcatgtttttttacagCAACAACGGGAAGAAAACTCTCTCTGTGCGCATCGTCCGCCACGCCTTCGAAATCATCCACC TGATGACGGACAAGAACCCCATTCAGGTCTTCGTCAACGCTGTTGAGAATGGCGGCCCCCGTGAAGACAGCACTCGAATCGGATCTGCTGGTGTTGTCAGACGCCAGGCTGTTGATGTGTCTCCCCTCCGCAGAGTGAACCAGGCCATCTACCTCATCTGCACCGGCGCAAGgtcagtccctctgaaagaaactgtacagctcctcacagagcggcagcgcgtggggtgttgaggtgtgtgtcagttcgtttgctgcacagtcagctgttgagtagtcgtcgctcttcaagatccgtccggcgtttgcgagtctctgccccagtctttcgtgaagcgtagagtcaacggcatgggtggatgtttttcagGCTGGCGGCCTTCCGGAACATCAAGACCATCGCCGAGTGCCTTGCGGACGAGATCATGAACTGCGCCAAGGAGTCCTCCAACGCTTACGCGATAAAGAAGAAGGACGAAATCGAGCGTGTTGCGAAGGCAAACCGATAA
5. UTRs are not necessary to successfully identify gRNAs, and so the 5’ and 3’ UTR input boxes can be left empty for this example.
6. Copy-paste the following sequence into the “**Please input the coding sequence of your protein here (no UTRs)**” box: ATGGCGACCGAGGCGAAGCTTTTCGGCCGCTGGTCGTACGACGATGTCAACGTCAGCGACCTTTCGCTCGTTGACTACATCGCGGTCAAGGACAAGGCCTGCGTCTTCGTGCCGCACACTGCTGGACGCTACCAGAAGAAGAGATTCCGCAAGGCCATGTGCCCCATTGTCGAGCGTCTCGTCAACTCCATGATGATGCACGGCCGCAACAACGGGAAGAAAACTCTCTCTGTGCGCATCGTCCGCCACGCCTTCGAAATCATCCACCTGATGACGGACAAGAACCCCATTCAGGTCTTCGTCAACGCTGTTGAGAATGGCGGCCCCCGTGAAGACAGCACTCGAATCGGATCTGCTGGTGTTGTCAGACGCCAGGCTGTTGATGTGTCTCCCCTCCGCAGAGTGAACCAGGCCATCTACCTCATCTGCACCGGCGCAAGGCTGGCGGCCTTCCGGAACATCAAGACCATCGCCGAGTGCCTTGCGGACGAGATCATGAACTGCGCCAAGGAGTCCTCCAACGCTTACGCGATAAAGAAGAAGGACGAAATCGAGCGTGTTGCGAAGGCAAACCGATAA
7. Select “**NGG**” when asked to “**Specify nuclease protospacer adjacent motif (PAM)**”.
8. Initially testing **OPTION 1**: In the “**Input the residue position of this amino acid**” box type “**143**”.
9. In the “**Please specify the maximum guide distance (in nucleotides) from the amino acid**” box type “**30**”.
10. Then click the “**Run Option 1**” box. A pop-up box will appear informing you when “**Your guide RNAs have successfully been generated**”. For Windows-based systems, the output .csv file is located within the same directory as the CRISPR-TAPE.exe file. On macOS the output .csv file is located within the CRISPR-TAPE.app package itself. This is accessible by right-clicking on the application and selecting “Show Package Contents” => subdirectory “Contents” => subdirectory “Resources”. The output .csv file from this test of Option 1 will be named “test” (see step 3 of this walk-through). The output file should match that shown in Figure 2a of the associated manuscript (see **Citation** below).
11. To test **OPTION 2**, amend the file name (see step 3 above) to “**test02**”. If the filename is not changed, the new output file overwrites the old output file. There is no need to change the inputs currently present in any of the other boxes as each run mode (Option 1 or 2) executes independently.
12. In the “**Input your target amino acid single letter code**” box, type “**C**”. Then click “**Run Option 2**”. This will identify the closest suitable gRNA 5’ and 3’ of each cysteine within the input coding sequence.
13. The successful identification of gRNA will be announced, and can be located as described in step 10 above. The output file should match that shown in Figure 2b of the associated manuscript (see **Citation** below).

### Troubleshooting

- All inputs are case sensitive, and it is important to adhere to the requirements specified adjacent to each entry box.
- gRNAs will be incorrect if exonic bases are not capitalised and intronic/untranslated bases are not lowercase in the genomic loci input.
- The programme currently only recognises standard “A”, “T”, “C” and “G” base nomenclature, and will filter out any other characters.
- If no protospacer adjacent motif is specified, the programme will not run and no guide RNAs will be outputted.
- The programme will stop running and a pop-up prompt will open if the inputted protein coding sequence and the exon sequences from the inputted genomic loci do not match.
- Off target counts will display as “-1” if gRNAs are not found within the forward or reverse complement of the organism genome. If this occurs, ensure the genomic loci and genome originate from the same organism.
- If some amino acids of a given type are missing from the OPTION 2 output, this is because they cannot be targeted using the current information. If this is the case, add more upstream and downstream bases to the input requesting 100 bases upstream and downstream of the genomic loci.
- Crashes are likely the result of your system not meeting the minimum system requirements (see **System requirements** above). In our experience this only occurs when working with genome greater than one giga-base in size.

### Error/bug reporting

We encourage all users to report any bugs they discover via the contact form at http://www.laboratorychild.com/contact

### Citation

TBC

## Notes

http://www.laboratorychild.com/crispr-tape

